# Differential effects of neurodegenerative disorders on verbal creative idea generation and its cognitive cornerstones

**DOI:** 10.1101/2025.04.22.650134

**Authors:** Melody M.Y. Chan, John D. O’Sullivan, Robert Adam, Philip E. Mosley, Michael J. Breakspear, Gail A. Robinson

**Author notes:** **Correspondence to:** Prof Gail A. Robinson Neuropsychology Research Clinic, School of Psychology, The University of Queensland, Brisbane QLD 4072, Australia.

## Abstract

Clinical research has documented a heterogeneous effect of neurodegenerative disorders on creativity. Some disorders are associated with impaired creative performance, while others may preserve, or even enhance, it. However, the underlying cognitive mechanisms that give rise to the heterogeneous behavioural manifestations remain poorly understood. From a theory-driven approach, we conducted a neuropsychological investigation to examine how frontotemporal lobar degeneration (FTLD), and Alzheimer’s Disease (AD) differentially affected the cognitive mechanisms underlying creative thought (semantic cognition, episodic memory, executive control functions). In parallel, we assessed participants’ verbal creative idea generation performance using the ideational fluency task (also known as the Alternate Uses Task). We analysed data from 24 individuals with FTLD [clinical consensus diagnosis corticobasal syndrome (CBS: n=20) or progressive supranuclear palsy (PSP: n=4)], 24 individuals with clinical consensus diagnosis AD, and 24 healthy individuals. We reached four major conclusions. First, while both disease groups exhibited deficits in controlled semantic retrieval and executive control functions, only participants with AD exhibited degradation in semantic representations and episodic memory. Second, both disease groups generated fewer responses in both typical and creative conditions across objects that were *context-free* - i.e., associated with multiple contexts (e.g., brick) and *context-bound* -i.e., associated with a dominant context (e.g., table knife). Third, when given a context-free object prompt, the ability to generate novel responses appeared to be preserved in participants with FTLD, which contrasted with an impaired ability when prompted to give creative uses for the context-bound object. Fourth, participants with AD produced similar levels of response novelty for both context-free and context-bound objects. From a clinical-cognitive neuroscience perspective, this study demonstrates that distinct cognitive profiles resulting from different neurodegenerative conditions yield differential effects on creative thought. As the first study to show that the contextual information of semantic stimuli can mediate the novelty of verbal responses, we suggest that future research should carefully examine how this factor influences creative task performance.

## Introduction

Creative thought is an intrinsic capacity that enables humans to generate novel and adaptive behaviours across many aspects of life. It is fundamental to scientific discoveries, art creations, and everyday problem solving. Like many other cognitive functions, creative thought is also affected by neurodegenerative diseases. Interestingly, this effect appears to be heterogeneous – some diseases lead to impaired performance in particular forms of creative mental activities, while others may preserve, or even enhance it. For instance, people with Alzheimer’s disease (AD) are found to have impaired verbal creativity (Marsh et al., 2024). In contrast, preserved creativity (Geser et al., 2021) or an even more fascinating cognitive phenomenon, the “release” of visuospatial creativity, has been reported in some individuals with frontotemporal lobar degeneration [FTLD; Erkkinen et al. (2018); Friedberg et al. (2023)]. Which cognitive mechanisms underpinning creative thought are affected by the diseases, and how the disease-induced cognitive changes affect creative mental activities remain largely unanswered.

Creativity has been predominantly viewed as a distinct mental activity that is associated with some domain-specific cognitive processes supporting flexible manipulation of acquired knowledge [e.g., divergent thinking (Guilford, 1967); associative thinking (Beaty & Kenett, 2023); conceptual expansion (Abraham et al., 2018)]. This view of creativity paves the way towards a more precise specification of the fundamental cognitive components underpinning creative thought (e.g., what are the knowledge sources, which processes support flexible manipulation of knowledge). To address and refine our understanding of creative thought, we have proposed a new conceptual framework for the cognitive and neural bases of creative thought by integrating not only the creativity research findings, but the broader clinical-cognitive neuroscience literature [i.e., *Cognitive Cornerstones Hypothesis*, Chan et al. (2023)]. Based on converging evidence (Chan et al., 2025a; Chan et al., 2025b), this hypothesis posits that creative thought comprises two cognitive cornerstones, *preexisting knowledge* and *executive mechanisms*. *Preexisting knowledge* refers to the reservoir from which ideas are drawn and synthesised. *Executive mechanisms* allow goal-directed, flexible, and transformative use of *preexisting knowledge*. Like many other aspects of higher cognition, creative thought arises from general purpose cognitive mechanisms supporting *semantic cognition*, *episodic memory*, and *executive control functions. Semantic cognition*, *episodic memory* and an intermediate mechanism that drives the integration and manipulation of buffered memories [which is often termed ‘working memory’, Baddeley, Baddeley (2000)] are the components of the *preexisting knowledge* cornerstone. Meanwhile, *energization*, *task-setting*, and *monitoring* (Stuss, 2011) are the components of the *executive mechanisms* cornerstone. The dynamic interplay of these cognitive mechanisms gives rise to a wide array of creative mental activities.

Importantly, FTLD and AD are known to differentially affect these cognitive mechanisms. For example, semantic cognition and episodic memory are disproportionately impacted in semantic dementia (a variant of FTLD) and AD, respectively (Nestor et al., 2006), while executive control deficits appear to be more generalised and are observed in both types of dementia (Burrell et al., 2014). If semantic cognition, episodic memory, and executive control functions underlie the manifestation of creative thought, then the differential impact of AD and FTLD on these cognitive domains would result in a different pattern of interplay between these mechanisms; such differences in cognitive profiles may explain the contrasting observations in creativity task performance in neurodegenerative diseases (e.g., Erkkinen et al., 2018; Friedberg et al., 2023; Geser et al., 2021; Marsh et al., 2024). In this neuropsychological investigation, we examined semantic cognition, episodic memory, and executive functions in individuals with FTLD, AD, and healthy counterparts. In parallel, we examined their verbal creative ability on an ideational fluency task (Lezak, 2012; also known as the Alternate Uses Task, AUT), a widely used task in creativity research that measures divergent thinking (i.e., the ability to generate multiple novel ideas). By exploring the cognitive cornerstones of creative thought alongside verbal creativity, this parallel investigation aims to reveal how disease-induced variations in cognitive abilities are linked to differences in creativity task performance. We hypothesise that FTLD and AD differentially impact the components of cognitive cornerstones (i.e., semantic cognition, episodic memory, executive control functions), which may lead to diverse cognitive phenomena in creative thought.

## Materials and methods

### Participants and sample size calculations

Included in this investigation were 72 native English speakers (aged 51-80) with normal/corrected-to-normal vision and hearing. This sample comprised three groups (**Table 1**) – an FTLD group [with patients diagnosed with Corticobasal Syndrome (CBS) or Progressive Supranuclear Palsy (PSP)], an AD group, and a healthy control group. The groups were matched by crystalised intelligence as a proxy for premorbid optimal level of function [measured by the National Adult Reading Test – 2nd edition (NART), (Bright et al., 2018)], biological sex, and handedness. This sample size was estimated based on Marsh et al. (2024), who reported a large effect size (Cohen’s d = 1.52, 95% CI [0.92, 2.12]) in verbal creativity impairment in AD patients when compared to healthy controls. Using the lower bound of this effect size, we determined that, for a power of 0.9 (with three groups and two covariates), 64 participants would be required to detect a statistically significant difference in verbal creativity among the three groups at α = .05. Participants were recruited for three separate research projects, each of which has its own ethics approval and specific participant inclusion criteria, as outlined below.

**Table 1:** Participants’ details.

| **Demographics** | **Healthy (n=24)** | **AD (n=24)** | **FTLD – CBS/PSP**<br>**(n=24)** | **F/$\chi^2$** |
| --- | --- | --- | --- | --- |
| **Age** | 68.96 (6.03) | 63.13 (7.63) | 68.88 (7.65) | 5.26** |
| **Premorbid crystallised intelligence** | 110.04 (5.90) | 104.38 (12.02) | 104.54 (9.43) | 3.87 |
| **Education level** | 15.00 (3.91) | 13.71 (3.56) | 12.20 (2.89) | 2.79* |
| **Sex (M:F)** | 8:16 | 7:17 | 10:14 | .86 |
| **Handedness (R:L)** | 23:1 | 22:2 | 23:1 | .53 |

Note:
Group comparison for scale data is performed with one-way ANOVA.
Group comparison for dichotomous data was performed with Chi-square test.
Significance levels:
\*p<.05
\*\*p<.01

Participants with FTLD (probable CBS n=20; probable PSP n=4) were recruited from the Royal Brisbane and Women’s Hospital (RBWH) Movement Disorder Clinic. According to relevant clinical diagnostic criteria [CBS: Mathew et al., 2012; PSP: Höglinger et al. (2017)], the CBS/PSP clinical diagnoses were made by consensus of the RBWH specialist movement disorder team led by consultant neurologists (J.O.S., R.A.). Participants’ core clinical features and principal diagnoses are summarised in **Table S1**. To ensure our observations were not confounded by motor or orofacial apraxia, we only included participants who did not show signs of apraxia on the Western Aphasia Battery-Revised (Kertesz, 2007) Apraxia subtest (median = 58; inter-quartile range = 56 – 60). This project was approved by the RBWH Human Research Ethics Committee (HREC; HREC/11/QRBWH/125) and The University of Queensland (UQ) HREC (2011000187).

Participants with probable AD (n=24) were recruited from the Prospective Imaging Study of Ageing: Genes, Brain and Behaviour (PISA) project (Lupton et al., 2021). Participants’ AD diagnosis was established during a multidisciplinary clinical consensus meeting (R.A., P.M., M.B., G.R.) based on criteria for AD (McKhann et al., 2011). Participants’ core clinical features and principal diagnoses are summarised in **Table S2**. This project was approved by the HRECs of QIMR Berghofer Medical Research Institute and RBWH (HREC/16/QRBWH/104) and UQ (2011000187).

Healthy controls (n=24) were recruited from the “Cognitive Functions and Neural Correlates” project approved by the UQ HREC (2011000187). This project was led by a consultant neuropsychologist (G.R.). Healthy controls were defined as individuals that were free from any medical, neurological or psychiatric conditions based on self-report, intact scores of 26 or above on the Montreal Cognitive Assessment [MoCA; Nasreddine et al. (2005)], and achieved scores above the 5^th^ percentile clinical cut-off in all cognitive domains assessed using a comprehensive neuropsychological battery, which was administered by clinical neuropsychology trainees (S.P. and C.G.) supervised by G.R..

### Procedures

All procedures were conducted in a quiet assessment room at the University of Queensland (Neuropsychology Research Clinic or School of Psychology) or on occasion at the participant’s home. Before participating in the study, all individuals provided written informed consent. A neuropsychological test battery was then conducted to assess performance across various cognitive domains, including attention, memory, executive control functions, language and novel idea generation (duration: approximately 3 hours). Participants were permitted to take breaks at any time as needed, without time restrictions. For all tests, the examiners gave clear instructions and examples before the testing phase, making sure participants were not under-performing due to misunderstanding of the instructions.

### Measures

Relevant to this investigation are standardised neuropsychological tests assessing the cognitive cornerstone components of creative thought (i.e., semantic cognition, episodic memory, and executive control functions) and a novel idea generation task tapping verbal creativity.

#### Preexisting knowledge

*Semantic cognition* was measured using the Graded Naming Test [GNT; McKenna and Warrington (1980); Warrington (1997)] and the semantic fluency test [1-minute, category: animals; Tombaugh et al. (1999)]. A higher number of correct responses on the GNT indicates a stronger ability to access and retrieve the meanings and names of words (i.e., *semantic representation*), and more responses generated on the semantic fluency test indicates a stronger ability in selectively retrieving relevant concepts from semantic memory (i.e., *semantic control*). *Episodic memory* was assessed using recognition trials from either the Warrington Recognition Memory Test for words (Warrington, 1984) or the Rey Auditory Verbal Learning Test – [RAVLT; Schmidt (1996)]. A greater hit rate (out of 100%) indicates a stronger ability to retrieve learned verbal information in the presence of distractors. The ability to *manipulate temporarily stored information* was measured using the Digit Span Backwards (DSB) subtest from the Wechsler Adult Intelligence Scale – 3rd Edition (Wechsler, 2000), with higher raw scores indicating better performance in manipulating buffered information.

#### Executive mechanisms

Different executive control processes (i.e., energization, task-setting, and monitoring) were evaluated using the phonemic fluency test (1-minute; Benton, 1968) and the colour-word Stroop test (Trenerry et al., 1989; Troyer et al., 2006). The phonemic fluency test requires participants to selectively retrieve words (but not proper nouns, numbers, or repeating words) that start with a specific letter (‘S’). By eliminating the semantic element, this test is thought to be a purer measure of verbal response initiation (Martin et al., 2021). The Stroop test requires participants to name the colour of the ink in which a word is printed as fast as they can, while the word itself names a different colour (i.e., the word ‘blue’ printed in red ink - incongruent condition). Performance is compared to a neutral word-reading condition, in which they read the word that matches the colour of the ink (i.e., ‘blue’ printed in blue ink - congruent condition). A higher proportion of responses generated in the first 15 seconds relative to the last 45 seconds of the phonemic fluency test reflects participants’ ability to initiate and sustain verbal responses (i.e., *energization:* Stuss et al., 1998). A smaller rate difference on the Stroop test, calculated as words per second in the incongruent condition minus that in the congruent condition, indicates better ability to select appropriate responses from competing alternatives (i.e., *task-setting*) while also *monitoring* responses over time and adjusting behaviour accordingly (Stuss and Alexander, 2007).

#### Creativity

The ideational fluency test (Lezak, 2012; Robinson et al., 2012, 2021) was used to assess novel idea generation. Participants were verbally prompted to generate as many uses as possible for two daily objects (i.e., “brick” and “table knife”) within 90 seconds under two conditions: typical uses or creative uses. The word prompt “brick” is context-free. In other words, it is associated with multiple contexts. For example, some participants may associate “brick” with “a rectangular block made of clay for construction”. For other participants, the word “brick” may evoke the mental representation of a “Lego brick”, which is made of plastic and used for children’s entertainment, or a type of “yoga block”, which can be made of wood or foam. In contrast, the prompt “table knife” is context-bound. The inclusion of the word “table” narrows its interpretation to a functional role within a dining scenario. This confines its attributes to those typically associated with dining utensils (e.g., made of stainless steel, paired with a fork). It is unlikely that participants would associate the term with other mental representations (e.g., a sharp knife designed for self-defence). Instead of single-word responses, participants often generate short phrases (e.g., “brick – for building a fence”).

Two measures are derived from this task to reflect participants’ verbal creativity performance: 1) the total number of valid responses generated in each condition for each object (i.e., response quantity), and 2) the averaged semantic distance between the object and the valid responses (i.e., response novelty). Responses were considered valid unless they were repeated within a condition or described object characteristics (e.g., “Brick – it is hard“; “Table knife – it is silverware”). The calculation of semantic distance involves two steps. First, the semantic textual similarity between the object prompt and each individual response was quantified using contextual embeddings generated by a pre-trained Sentence Transformer Model (SBERT *all-MiniLM-L6-v2*; (Reimers & Gurevych, 2019) that leverages large-scale natural language datasets. Representing textual similarity using sentence embeddings generated by pre-trained, context-specific large language models, but not context-independent additive or multiplicative vector models [e.g., Beaty and Johnson (2021)], allows more accurate numerical representation of the semantic similarity between stimuli and responses. Second, the semantic distance (i.e., 1 − *cosθ*) between the object prompt and each individual response was calculated using the following formula: 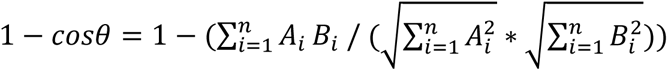. A larger 1 − *cosθ* value indicates greater semantic distance, which has been used as an indicator of novelty (Beaty & Johnson, 2021). These values were averaged for each condition to represent the novelty of responses generated. To ensure the robustness of semantic similarity measures derived from SBERT, which was originally designed for full-sentence inputs, we conducted parallel analyses using both the raw participant responses (typically short phrases) and syntactically rephrased, complete sentences. This allowed us to evaluate the consistency of semantic distance estimates across preprocessing formats. Participants’ data were processed with in-house Python script available at https://osf.io/2u5jv/.

### Statistical analysis

Normality of the raw data was checked with the Shapiro-Wilk test. As the three groups were not matched for age and education level, these factors were included as covariates in the factorial analyses.

To compare neuropsychological performance between the patient and control groups, a univariate ANCOVA (or Quade nonparametric ANCOVA for non-normally distributed data) was conducted on the raw data representing each cognitive domain: *semantic cognition* (GNT total correct and total number of responses generated in semantic fluency task), *episodic memory* (RMT – word/RAVLT recognition trials: hit rate), *manipulation of temporarily stored information* (DSB raw score), and *executive control functions* (total responses generated in the first 15 seconds of the phonemic fluency test, and the rate difference between incongruent and congruent condition in Stroop test). To control for an overall error rate at 5%, only a between-group difference with a significance level of p < .05/6 = .0083 (i.e., Bonferroni correction for alpha=.05) was considered statistically significant for each individual test component.

To examine the effect of different neurodegenerative disorders on creativity performance, a 3 (group: FTLD, AD, healthy control) x 2 (condition: typical, creative) x 2 (object: brick, table knife) mixed ANCOVA was used with age and education level as covariates. The dependent variable was the averaged semantic distance (i.e., response novelty) and the number of valid responses generated (i.e., response quantity). As most of these data were not normally distributed, Aligned Rank Transform (ART, Wobbrock et al, 2011) was performed on all data; ANCOVA was performed with the ART-transformed data. Significant interaction effects and main effects of the group were followed by post-hoc pairwise comparisons with Bonferroni corrections applied (ART-Contrast; Elkin et al., 2021).

To explore the relationship between the cognitive cornerstones components and creativity task performance, Spearman’s rho nonparametric correlation analysis was performed using the raw scores of the measures. Given the exploratory nature of these analyses, correlation coefficients that survived a False Discovery Rate (FDR) correction with p < .05 [Benjamini-Hochberg procedure; Benjamini and Hochberg (1995)] were considered statistically significant.

Normality check, factorial and correlation analyses were performed using SPSS Statistics 29 software (IBM, 2023). ART/ART-Contrast was performed using ARTool version 2.2.2.

## Results

### Cognitive cornerstones of creative thought

Participants’ performance on tasks tapping the cognitive cornerstones components are summarised in **Table 2**. Compared to the healthy control group, 1) both disease groups performed significantly worse on tasks tapping controlled semantic retrieval [post-hoc pairwise comparisons: AD < healthy (p<.001), CBS/PSP < healthy (p<.001)], energization [post-hoc pairwise comparisons: AD < healthy (p<.01), CBS/PSP < healthy (p<.001)], and task-setting/monitoring [post-hoc pairwise comparisons: AD < healthy (p<.01), CBS/PSP < healthy (p<.001)]; 2) semantic representations and episodic memory were only degraded in participants with AD [post-hoc pairwise comparisons: AD < healthy (all p <.001), AD < CBS/PSP (all p <.05)], but not those with FTLD (CBS/PSP < healthy n.s.); and 3) no significant differences were observed in tasks tapping the manipulation of temporarily stored information F_2,67_ = 2.15, p = .13).

**Table 2:**
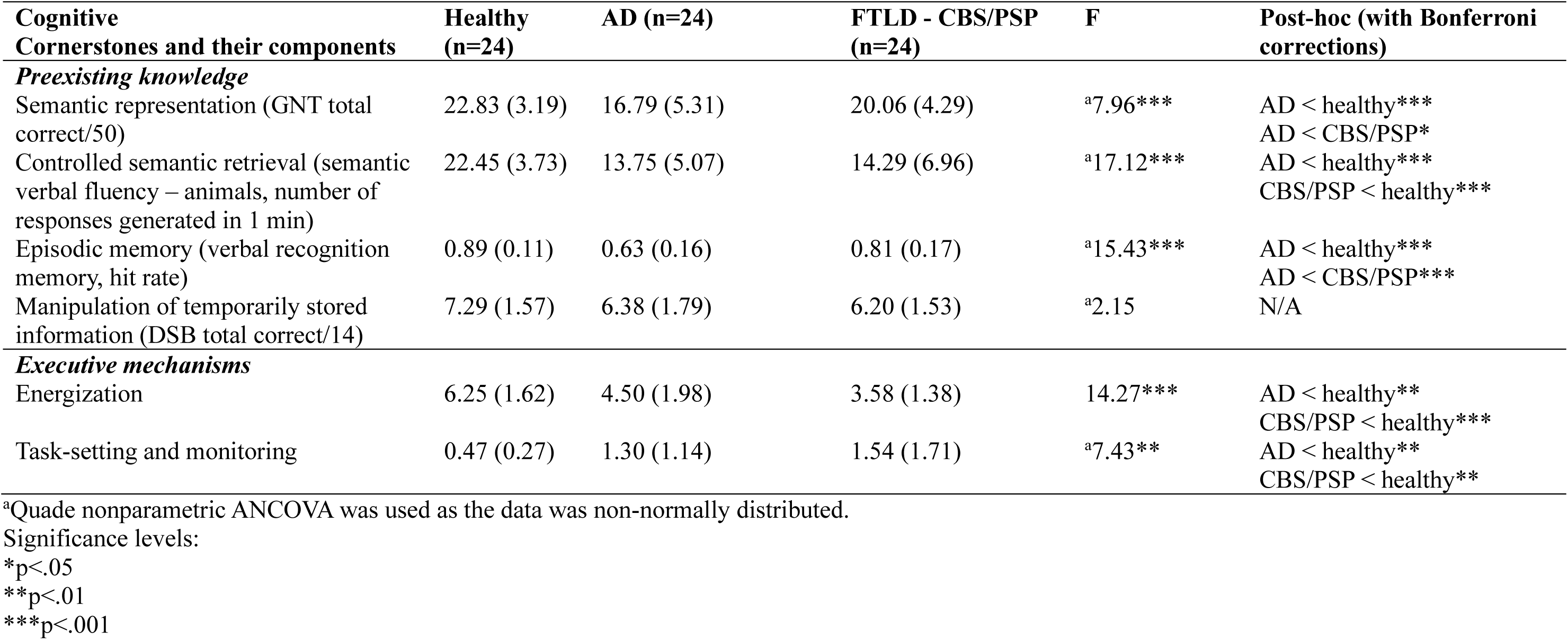
Neuropsychological performance.

| **Cognitive Cornerstones and their components** | **Healthy (n=24)** | **AD (n=24)** | **FTLD - CBS/PSP (n=24)** | **F** | **Post-hoc (with Bonferroni corrections)** |
| --- | --- | --- | --- | --- | --- |
| ***Preexisting knowledge*** |  |  |  |  |  |
| Semantic representation (GNT total correct/50) | 22.83 (3.19) | 16.79 (5.31) | 20.06 (4.29) | <sup>a</sup>7.96*** | AD < healthy***<br>AD < CBS/PSP* |
| Controlled semantic retrieval (semantic verbal fluency – animals, number of responses generated in 1 min) | 22.45 (3.73) | 13.75 (5.07) | 14.29 (6.96) | <sup>a</sup>17.12*** | AD < healthy***<br>CBS/PSP < healthy*** |
| Episodic memory (verbal recognition memory, hit rate) | 0.89 (0.11) | 0.63 (0.16) | 0.81 (0.17) | <sup>a</sup>15.43*** | AD < healthy***<br>AD < CBS/PSP*** |
| Manipulation of temporarily stored information (DSB total correct/14) | 7.29 (1.57) | 6.38 (1.79) | 6.20 (1.53) | <sup>a</sup>2.15 | N/A |
| ***Executive mechanisms*** |  |  |  |  |  |
| Energization | 6.25 (1.62) | 4.50 (1.98) | 3.58 (1.38) | 14.27*** | AD < healthy**<br>CBS/PSP < healthy*** |
| Task-setting and monitoring | 0.47 (0.27) | 1.30 (1.14) | 1.54 (1.71) | <sup>a</sup>7.43** | AD < healthy**<br>CBS/PSP < healthy** |

<sup>a</sup>Quade nonparametric ANCOVA was used as the data was non-normally distributed.
Significance levels:
\*p<.05
\*\*p<.01
\*\*\*p<.001

### Creativity task performance

Participants’ performance in the novel idea generation task is illustrated in **Figure 1**.

**Figure 1:**
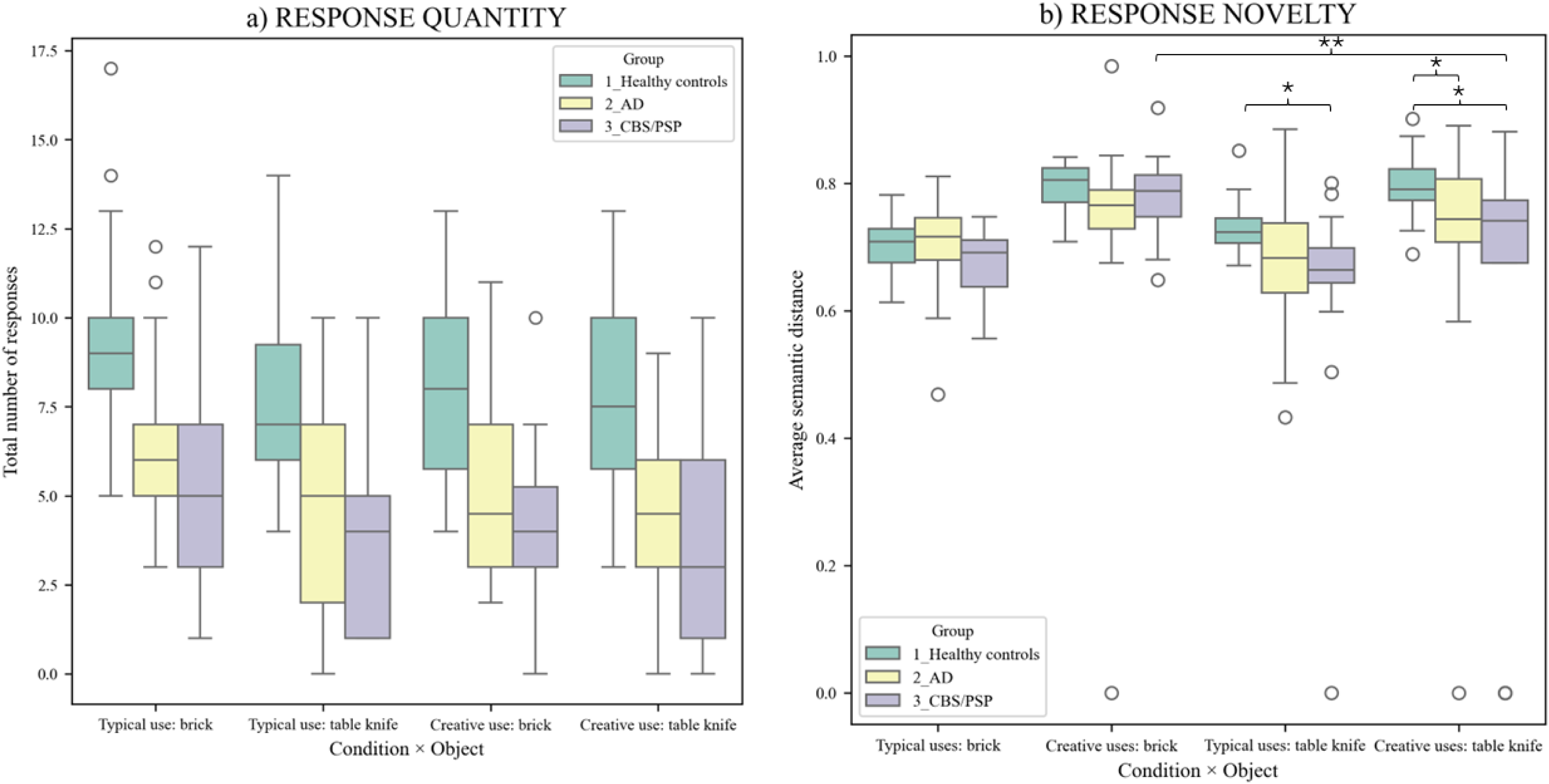
Novel idea generation task performance. Note: Colour of the bars: blue = healthy controls; red = AD patients; teal = FTLD patients (i.e., CBS/PSP) Significance level: *p<.05; **p<.01

#### Response quantity (*Figure 1a*)

Compared to the healthy control group, both disease groups generated fewer typical and creative responses for both objects, brick and table knife (main effect of group: F_2,67_ = 20.35, p<.001, ηp2 = .38; pairwise comparison AD < healthy p<.001; pairwise comparison CBS/PSP < healthy p < .001; pairwise comparison CBS/PAP and AD p = .93). Visual inspection of the **Fig. 1a** suggests that healthy controls tended to generate a lower number of typical uses for table knife than for brick, but post-hoc pairwise comparison revealed a nonsignificant difference (p = .57). The group*object*condition interaction effect was nonsignificant (F_2,67_ = 1.78, p = .18).

#### Response novelty (*Figure 1b*)

As analyses using ART-transformed data yielded converging findings across both raw responses and syntactically rephrased sentences, we report here the results based on raw responses. A significant group*object*condition interaction effect was observed (F_2,67_ = 4.92, p = .010, ηp2 = .13). This was driven by three observations. First, the patients generated much less novel responses for table knife [pairwise comparison (table knife – unconventional uses) AD < healthy p=.039; pairwise comparison CBS/PSP < healthy p = .010; pairwise comparison CBS/PSP and AD n.s.], but not for brick (pairwise comparisons all n.s.). Second, when prompted to generate typical uses of the table knife, CBS/PSP patients (p = .043), but not the AD patients (p = .095), generated responses with significantly lower novelty than the healthy controls. Third, the CBS/PSP patient group generated significantly more novel responses for the brick but less novel responses for the table knife in the creative condition (p = .006).

### Correlation analysis

The nonparametric correlation results are shown in **Table 3**. The results reveal the following: 1) better semantic representation was associated with the generation of more novel creative responses (ρ_71_ = .340, FDR-corrected p = .026) and more typical responses (ρ_71_ = .331, FDR-corrected p = .030) of the brick; 2) better controlled semantic retrieval was associated with the generation of more novel responses when asked to produce typical uses of the table knife, and with generating more responses overall (ρ_71_ > .418, all FDR-corrected p < .001); 3) better episodic memory was associated with the generation of more typical uses of the brick (ρ_71_ = .331, FDR-corrected p <.05); 4) better manipulation of buffered information (ρ_71_ = .298, FDR-corrected p <.05) and interference control (ρ_71_ = −.359, FDR-corrected p <.01) were associated with the generation of more novel creative uses for the table knife; and 5) better energization is associated with generating more responses overall (ρ_71_ > .393, all FDR-corrected p < .01).

**Table 3:** Correlation table.

| Bivariate correlation (rho) |  | Creativity task performance –<br>response quantity |  |  |  | Creativity task performance –<br>response novelty |  |  |  |
| --- | --- | --- | --- | --- | --- | --- | --- | --- | --- |
|  |  | *Object: Brick* |  | *Object: Table knife* |  | *Object: Brick* |  | *Object: Table knife* |  |
|  |  | Typical | Creative | Typical | Creative | Typical | Creative | Typical | Creative |
| Cognitive<br>cornerstones<br>components | Semantic representation | .331* | .289 | .160 | .268 | .074 | .340* | .245 | .056 |
|  | Controlled semantic retrieval | .510*** | .615*** | .471*** | .576*** | .002 | .236 | .418*** | .077 |
|  | Episodic memory | .338* | .259 | .132 | .306* | -.066 | .230 | .144 | .130 |
|  | Manipulation of temporarily stored information | .188 | .147 | .008 | .253 | -.062 | .137 | .270 | .298* |
|  | Initiation | .464*** | .393** | .387** | .403*** | -.059 | -.040 | .250 | .213 |
|  | Task-setting and monitoring | -.289 | -.249 | -.150 | -.296 | .133 | -.086 | -.233 | -.359** |

Significance levels (FDR-corrected):
\*p<.05; \*\*p<.01, \*\*\*p<.001

## Discussion

Through the lens of clinical-cognitive neuroscience, this study offers a new perspective to understand how creativity is compromised in the context of neurodegenerative disorders. Specifically, we examined how FTLD (CBS/PSP) and AD affect verbal creativity, and the general-purpose cognitive mechanisms underpinning creative thought (i.e., semantic cognition, episodic memory, executive control functions). The FTLD and AD groups neuropsychological performance was compared to healthy controls. Four major findings were revealed: 1) the disease groups showed partially overlapping patterns of impairment in the cognitive cornerstones of creative thought; 2) both disease groups generated fewer responses in both typical and creative conditions across both context-free (i.e., brick) and context-bound (i.e., table knife) objects; 3) both patient groups were able to generate novel responses when given a context-free object prompt (e.g., brick); however, FTLD patients generated significantly less novel uses for context-bound objects (e.g., table-knife); 4) participants with AD produced similar levels of response novelty for both context-free and context-bound objects.

Compared to their healthy counterparts, individuals with neurodegenerative diseases produced significantly fewer typical and creative uses of common objects. Correlation analysis further shows that a higher number of typical and creative uses generated is strongly associated with better performance on the semantic and phonemic fluency tasks. Our results imply that the efficiency with which one could verbally generate creative ideas depends on more domain-general cognitive processes supporting voluntary generation of non-overlearned responses (Robinson et al., 2012). These processes include controlled semantic retrieval and energization. Visual inspection of data revealed that healthy participants tended to respond differently when they were given different object prompts. For instance, an object prompt without a predefined context (i.e., brick) elicited a broader range of typical uses in healthy controls. This largely aligns with the concept of *semantic diversity*, which suggests that the meanings of words vary depending on their contexts (Hoffman et al., 2013). In contrast, individuals with neurodegenerative disorders generated comparable typical responses when given different object prompts. In other words, their ability to generate diverse typical responses is uniformly impaired regardless of context. Although our study lacked adequate power to detect within-group differences in the healthy control group, future large-scale studies could further explore this relationship by asking participants to generate both typical and creative uses for a broader range of objects, incorporating those with well-defined contexts as well as those that are context-free.

Although it is yet to be explored whether semantic diversity mediates creative performance, we are certain that contextual information mediates verbal creativity. Regarding the context-free object (i.e., brick), the responses that healthy controls generated were significantly more novel (indexed by greater averaged semantic distance between object prompt and responses) during the creative condition than the typical condition. Such difference between the creative and typical conditions disappeared for the context-bound object (i.e., table knife). This finding suggests that in healthy ageing the contextual information of semantic stimuli can mediate verbal creativity. For context-free objects like the brick, which have no fixed associations, the creative condition prompts individuals to explore a broader semantic space. The absence of preestablished associations enables healthy individuals to draw on more remotely associated concepts, which facilitates novel idea generation. The correlation results further justify our claim indicating that people with richer semantic representations tend to generate responses with greater novelty in the brick creative condition. Conversely, objects with strong associations with a particular context, like the table knife, impose inherent contextual constraints. To compare and contrast both dominant ideas (e.g., table knife is relatively blunt; it is kitchenware etc.) and nondominant representations (e.g., the blade of a table knife is flat and narrow, which resembles a flat-head screwdriver), one would rely on better temporary storage of retrieved knowledge in the multimodal knowledge buffer (to concurrently hold various representations for goal-directed evaluation) and the voluntary suppression of dominant ideas, which primarily involves task-setting and monitoring (Stuss, 2011). Our claim is again supported by the correlation results, which suggest that people with better ability to manipulate buffered knowledge and perform better in an interference control task (i.e., Stroop test) tend to achieve greater response novelty in the table knife creative condition.

The results from the patient groups align with our observations within the healthy control group. For instance, AD patients showed no difference in response novelty across objects (i.e., brick/table knife) and conditions (i.e., typical/creative uses), which reflects generalised impairments in semantic and episodic memories. Degraded knowledge sources likely limit the diversity of ideas they can draw from for both context-free and context-bound objects. In contrast, patients with FTLD exhibited preserved creative performance for the brick (comparable to healthy controls), which could be explained by their intact semantic representations and episodic memory. However, their creative performance significantly declined for the table knife, highlighting impairments in energization, working memory, and interference control, which is critical for suppressing dominant associations. These findings highlight the dynamic contributions of the cognitive cornerstones to verbal creativity, which is mediated by task demands (i.e., creative versus typical and context-free versus context-bound), and how verbal creativity is preserved, or compromised, in the context of neurodegenerative disorders. Future research should explore how our brain supports the complex interactions between the cognitive cornerstones under different task demands, and how that would change when some brain functions are affected due to neurodegenerative diseases. It would also be interesting to explore whether our observations in the clinical groups translates to nonverbal creativity tasks.

Creative thought has historically been studied in isolation from broader clinical-cognitive neuroscience. As a result, the cognitive mechanisms underpinning creative thought remain underspecified, making it challenging to understand why creativity is compromised, preserved, or enhanced in the context of neurological disorders. This study serves as a worked example of how creative thought can be understood within a neurocognitive framework, linking general purpose cognitive mechanisms related to creative thought (i.e., semantic cognition, episodic memory, executive control functions), to creativity task performance associated with different neurodegenerative diseases. By focusing on how major cognitive processes supporting flexible manipulation of knowledge influences verbal creative idea generation, we move beyond studying creativity phenomena in isolation, grounding observations in a theoretical framework. Despite its important contribution to the field, we acknowledge the potential for underlying pathological heterogeneity in clinically defined CBS. While previous works suggest that CBS can arise from underlying AD pathology, this is unlikely in our cohort. The clinical profiles of our CBS patients were characterised by prominent asymmetric motor and cortical signs (e.g., apraxia, dystonia, alien limb phenomenon), rather than early memory or visuospatial deficits typically seen in CBS with underlying AD pathology (Wilson et al., 2021). In addition, we compared the CBS group against a separate cohort with well-characterised AD (including amyloid positive on PET imaging), which helps minimise diagnostic overlap. Future experimental research should clarify how contextual information affects creative idea generation and explore its neural correlates. Moreover, clinical research should explore the specific mechanisms driving the release of creativity in some individuals with FTD, with a focus on visuospatial creativity tasks.

This study highlights the dynamic interplay between general purpose cognitive mechanisms and object properties in shaping verbal creativity. Furthermore, it can be viewed as a worked example of how to apply the “*Cognitive Cornerstones Hypothesis*” framework to understand creative thought; that is, when viewed through the lens of clinical-cognitive neuroscience. Broadly speaking, our findings support this Hypothesis: 1) the ability to generate multiple novel responses relies on general purpose cognitive mechanisms supporting semantic cognition, episodic memory, and executive control mechanisms; and 2) the dynamic interplay between these mechanisms supports different aspects of verbal creativity (i.e., response quantity and response novelty). Using disease models, we demonstrated that differential impairments in these mechanisms due to neurodegenerative disorders lead to distinct effects on verbal creativity performance. Future research should build on these findings by investigating how object use diversity mediates creative response generation, exploring the dynamic interaction of cognitive cornerstones in nonverbal creativity, and understanding how these processes are compromised in the context of neurodegenerative diseases.

## Data availability

Anonymised dataset used in this study will be available upon request.

## Acknowledgements

The authors would like to thank Ms. Amelia Ceslis for assistance with collecting the AD data and Ms. Casey Gilbert and Ms. Samara Phelan for their efforts in assisting with the recruitment of healthy controls, and the participants with AD/CBS/PSP who enrolled in the studies.

---

**Table S1:** Core clinical features and principal diagnoses of participants with primary tauopathies.

| **Subject code** | **Core clinical features** | **Principal diagnosis** |
| --- | --- | --- |
| CBD1 | Insidious onset, no response to levodopa, limb apraxia, speech impairment – dysarthria, right food dystonia, cortical sensory loss (dystereognosis) | Probable CBS |
| CBD2 | Insidious onset, no response to levodopa, akinetic rigid syndrome, limb apraxia, focal/segmental myoclonus, frontal executive dysfunction | Probable CBS |
| CBD3 | Insidious onset, no sustained response to levodopa, akinetic rigid syndrome, focal or segmental myoclonus, asymmetric dystonia (upper limb), alien limb | Probable CBS |
| CBD4 | Insidious onset, no sustained response to levodopa, limb apraxia, asymmetrical dystonia, frontal executive dysfunction | Probable CBS |
| CBD5 | Insidious onset, no sustained response to levodopa, akinetic rigid syndrome, speech and language impairment (dysgraphia), focal/segmental myoclonus, asymmetrical dystonia | Probable CBS |
| CBD6 | Marked apraxia in both upper and lower limbs, worse on the right; no orobuccal apraxia. Rigidity in the right upper limb with mild intention tremor. No resting tremor. Dystonic posturing of the right foot particularly while walking. No cortical sensory loss. No gaze limitation. Speech normal. | Probable CBS |
| CBD7 | Insidious onset, no sustained response to levodopa, limb apraxia (R>L), speech and language impairment, frontal executive dysfunction | Probable CBS |
| CBD8 | Speech and language impairment (dysgraphia), right lower limb dyskinesia | Probable CBS |
| CBD9 | Right limb dyskinesia, speech and language impairment (effortful speech) | Probable CBS |
| CBD10 | Insidious onset, limb apraxia, akinetic rigid syndrome, speech and language impairment (aphasia), frontal executive dysfunction | Probable CBS |
| CBD11 | Insidious onset, no sustained response to levodopa, akinetic rigid syndrome, limb apraxia, dysgraphaesthesia | Probable CBS |
| CBD12 | Insidious onset, no sustained response to levodopa, akinetic rigid syndrome, speech and language impairment – mild dysarthria | Probable CBS |
| CBD13 | Insidious onset, right lower limb apraxia, no response to levodopa, speech and language impairment (effortful speech) | Probable CBS |
| CBD14 | Insidious onset, no sustained response to levodopa, akinetic rigid syndrome, speech and language impairment (dysarthria), frontal executive dysfunction (disinhibition) | Probable CBS |
| CBD15 | Moderate dyspraxia worse in the left than the right limbs; dyspraxic gait; mild dysarthria but no orobuccal dyspraxia; no cortical sensory signs, lack of response to levodopa | Probable CBS |
| CBD16 | Insidious onset, limb apraxia, speech and language impairment (dysgraphia), executive dysfunction, behavioural change (apathy, stereotyped phrases, flat affect) | Probable CBS |
| CBD17 | Insidious onset, no response to levodopa, asymmetrical dystonia, speech and language impairment (dysgraphia), limb apraxia | Probable CBS |
| CBD18 | Insidious onset, no sustained response to levodopa, akinetic rigid syndrome, asymmetrical dystonia | Probable CBS |
| CBD19 | Insidious onset, no sustained response to levodopa, akinetic rigid syndrome, limb apraxia | Probable CBS |
| CBD20 | Insidious onset, no sustained response to levodopa, akinetic rigid syndrome, alien limb | Probable CBS |
| PSP1 | Reduced blink rate with mild frontalis overactivity, restricted range of vertical up and down gaze with jerky horizontal and vertical pursuit, delayed and slow saccadic eye movements with up gaze worse than down gaze which was worse than horizontal gaze; 5 unprovoked falls within 3 years. | Probable PSP |
| PSP2 | Vertical gaze palsy ++, speech and language disorders | Probable PSP |
| PSP3 | Slowed vertical saccades, averaged speech, impaired postural reflexes, mild shuffle, mild bilateral bradykinesia, no proximal or distal rigidity. Repeated unprovoked falls within 3 years | Probable PSP |
| PSP4 | Restricted up gaze and definite slowing of horizontal and vertical upward saccadic eye movements. Repeated unprovoked falls, frequently occur when being stationary rather than walking. | Probable PSP |

Note: All patients had a minimum duration of symptoms > 1 year and were > 50 years.

**Table S2:** Core clinical features and principal diagnosis of participants with secondary tauopathy.

| **Subject code** | **Diagnostic criterion A2: cognitive domains impaired (according to standardised neuropsychological testing<sup>a</sup>)** | **Principal diagnosis (variant)** |
| --- | --- | --- |
| PISA1 | Memory, Language, Executive, Attention | AD (Amnestic) |
| PISA2 | Memory, Language | AD (Amnestic) |
| PISA3 | Memory, Numeracy, Executive, Attention | AD (Amnestic) |
| PISA4 | Memory, Language, Attention | AD (Amnestic) |
| PISA5 | Memory, Numeracy, Executive, Attention, Speed | AD (Executive) |
| PISA6 | Numeracy, Attention, Speed | AD (Visual) |
| PISA7 | Memory, Numeracy, Executive, Attention, Speed | AD (Amnestic) |
| PISA8 | Memory, Language, Literacy, Numeracy, Executive, Attention, Speed | AD (Amnestic) |
| PISA9 | Memory, Executive | AD (Amnestic) |
| PISA10 | Memory, Language, Executive, Attention | AD (Executive) |
| PISA11 | Memory, Numeracy, Executive, Speed | AD (Amnestic) |
| PISA12 | Memory, Numeracy, Executive, Attention, Speed | AD (Amnestic) |
| PISA13 | Memory, Executive | AD (Amnestic) |
| PISA14 | Memory, Numeracy, Executive, Attention, Speed | AD (Amnestic) |
| PISA15 | Memory, Language, Literacy, Numeracy, Executive, Attention | AD (Visual) |
| PISA16 | Language, Numeracy, Executive, Attention | AD (Language) |
| PISA17 | Executive, Attention, Speed | AD (Visual) |
| PISA18 | Memory, Executive, Attention, Speed | AD (Amnestic) |
| PISA19 | Memory, Executive | AD (Amnestic) |
| PISA20 | Memory, Numeracy, Executive, Speed | AD (Amnestic) |
| PISA21 | Memory, Executive, Attention, Speed | AD (Amnestic) |
| PISA22 | Memory, Language | AD (Amnestic) |
| PISA23 | Memory, Numeracy, Executive, Attention, Speed | AD (Visual) |
| PISA24 | Memory, Executive, Attention | AD (Amnestic) |

Note: For all patients, significant cognitive decline was reported by a knowledgeable informant (criterion A1); these cognitive deficits interfered with independence in instrumental activities of daily living (criterion B); these cognitive deficits did not occur exclusively in the context of delirium (criterion C), and these deficits were not better explained by another mental disorder (criterion D).
<sup>a</sup> For the tests used, please see Lupton et al. (2021).

## References

Abraham, A., Rutter, B., Bantin, T., & Hermann, C. (2018). Creative conceptual expansion: A combined fMRI replication and extension study to examine individual differences in creativity. Neuropsychologia, 118, 29–39.

American Psychiatric Association, D. (2013). Diagnostic and statistical manual of mental disorders: DSM-5 (Vol. 5). American psychiatric association Washington, DC.

Baddeley, A. (2000). The episodic buffer: a new component of working memory? Trends in cognitive sciences, 4(11), 417–423.

Beaty, R. E., & Johnson, D. R. (2021). Automating creativity assessment with SemDis: An open platform for computing semantic distance. Behavior research methods, 53(2), 757–780.

Beaty, R. E., & Kenett, Y. N. (2023). Associative thinking at the core of creativity. Trends in cognitive sciences.

Benjamini, Y., & Hochberg, Y. (1995). Controlling the false discovery rate: a practical and powerful approach to multiple testing. Journal of the Royal statistical society: series B (Methodological), 57(1), 289–300.

Bright, P., Hale, E., Gooch, V. J., Myhill, T., & van der Linde, I. (2018). The National Adult Reading Test: restandardisation against the Wechsler adult intelligence scale—fourth edition. Neuropsychological Rehabilitation, 28(6), 1019–1027.

Burrell, J. R., Hodges, J. R., & Rowe, J. B. (2014). Cognition in corticobasal syndrome and progressive supranuclear palsy: a review. Movement Disorders, 29(5), 684–693.

Chan, M. M., Lambon Ralph, M., & Robinson, G. A. (2025a). The neural basis of creative thought: An activation likelihood estimation meta-analysis involving over 17,000 participants. bioRxiv, 2025–02.

Chan, M. M., Cho, E., Lambon Ralph, M., & Robinson, G. A. (2025b). The cognitive and neural bases of creative thought: a cross-domain meta-analysis of transcranial direct current stimulation studies. bioRxiv, 2025–03.

Chan, M. M., Lambon Ralph, M., & Robinson, G. (2023). The neurocognitive cornerstones of creative thought. 10.31234/osf.io/mv2q7

Chung, D.-e. C., Roemer, S., Petrucelli, L., & Dickson, D. W. (2021). Cellular and pathological heterogeneity of primary tauopathies. Molecular neurodegeneration, 16, 1–20.

Dabul, B. L. (2000). ABA-2: Apraxia Battery for Adults-Second Edition.

Elkin, L. A., Kay, M., Higgins, J. J., & Wobbrock, J. O. (2021, October). An aligned rank transform procedure for multifactor contrast tests. In The 34th annual ACM symposium on user interface software and technology (pp. 754–768).

Erkkinen, M. G., Zúñiga, R. G., Pardo, C. C., Miller, B. L., & Miller, Z. A. (2018). Artistic renaissance in frontotemporal dementia. Jama, 319(13), 1304–1306.

Field, A. (2024). Discovering statistics using IBM SPSS statistics. Sage publications limited.

Friedberg, A., Pasquini, L., Diggs, R., Glaubitz, E. A., Lopez, L., Illán-Gala, I., Iaccarino, L., La Joie, R., Mundada, N., & Knudtson, M. (2023). Prevalence, timing, and network localization of emergent visual creativity in frontotemporal dementia. JAMA neurology, 80(4), 377–387.

Geser, F., Jellinger, K. A., Fellner, L., Wenning, G. K., Yilmazer-Hanke, D., & Haybaeck, J. (2021). Emergent creativity in frontotemporal dementia. Journal of Neural Transmission, 128, 279–293.

Guilford, J. P. (1967). Creativity: Yesterday, today and tomorrow. The Journal of Creative Behavior, 1(1), 3–14.

Hoffman, P., Lambon Ralph, M. A., & Rogers, T. T. (2013). Semantic diversity: A measure of semantic ambiguity based on variability in the contextual usage of words. Behavior research methods, 45, 718–730.

Höglinger, G. U., Respondek, G., Stamelou, M., Kurz, C., Josephs, K. A., Lang, A. E., Mollenhauer, B., Müller, U., Nilsson, C., & Whitwell, J. L. (2017). Clinical diagnosis of progressive supranuclear palsy: the movement disorder society criteria. Movement disorders, 32(6), 853–864.

Hutchinson, A., & Mathias, J. L. (2007). Neuropsychological deficits in frontotemporal dementia and Alzheimer’s disease: a meta-analytic review. *Journal of Neurology*, Neurosurgery & Psychiatry, 78(9), 917–928.

Josephs, K. A., Knopman, D., Whitwell, J., Boeve, B., Parisi, J., Petersen, R., & Dickson, D. (2005). Survival in two variants of tau-negative frontotemporal lobar degeneration: FTLD-U vs FTLD-MND. Neurology, 65(4), 645–647.

Kovacs, G. G. (2015). Invited review: neuropathology of tauopathies: principles and practice. Neuropathology and applied neurobiology, 41(1), 3–23.

Lezak, M. D. (2012). Neuropsychological assessment. Oxford University Press, USA.

Lupton, M. K., Robinson, G. A., Adam, R. J., Rose, S., Byrne, G. J., Salvado, O., Pachana, N. A., Almeida, O. P., McAloney, K., & Gordon, S. D. (2021). A prospective cohort study of prodromal Alzheimer’s disease: prospective imaging study of ageing: genes, brain and behaviour (PISA). NeuroImage: Clinical, 29, 102527.

Marsh, G., Aung, O., Ceslis, A., Adam, R., Mosley, P., Fripp, J., & Robinson, G. (2024). Generation of novel ideas: Creativity in Alzheimer’s disease, mild cognitive impairment, and healthy older adults. Creativity research journal. 10.1080/10400419.2024.2304498

Martin, A. K., Barker, M., Gibson, E., & Robinson, G. (2021). Response initiation and inhibition and the relationship with fluid intelligence across the adult lifespan. Archives of clinical neuropsychology, 36(2), 231–242.

Mathew, R., Bak, T. H., & Hodges, J. R. (2012). Diagnostic criteria for corticobasal syndrome: a comparative study. *Journal of Neurology*, Neurosurgery & Psychiatry, 83(4), 405–410.

McKenna, P., & Warrington, E. K. (1980). Testing for nominal dysphasia. *Journal of Neurology*, Neurosurgery & Psychiatry, 43(9), 781–788.

Nasreddine, Z. S., Phillips, N. A., Bédirian, V., Charbonneau, S., Whitehead, V., Collin, I., Cummings, J. L., & Chertkow, H. (2005). The Montreal Cognitive Assessment, MoCA: a brief screening tool for mild cognitive impairment. Journal of the American Geriatrics Society, 53(4), 695–699.

Nestor, P. J., Fryer, T. D., & Hodges, J. R. (2006). Declarative memory impairments in Alzheimer’s disease and semantic dementia. NeuroImage, 30(3), 1010–1020.

Reimers, N., & Gurevych, I. (2019). Sentence-BERT: Sentence Embeddings using Siamese BERT-Networks. arXiv preprint arXiv:1908.10084.

Robinson, G., Shallice, T., Bozzali, M., & Cipolotti, L. (2012). The differing roles of the frontal cortex in fluency tests. Brain, 135(7), 2202–2214.

Schmidt, M. (1996). Rey Auditory verbal learning test: A handbook. Western Psychological Services.

Stuss, D. T. (2011). Functions of the frontal lobes: relation to executive functions. Journal of the international neuropsychological Society, 17(5), 759–765.

Tombaugh, T. N., Kozak, J., & Rees, L. (1999). Normative data stratified by age and education for two measures of verbal fluency: FAS and animal naming. Archives of clinical neuropsychology, 14(2), 167–177.

Trenerry, M., Crosson, B., DeBoe, J., & Leber, W. (1989). Stroop Neuropsychological Screening Test manual. Psychological Assessment Resources.

Troyer, A. K., Leach, L., & Strauss, E. (2006). Aging and response inhibition: Normative data for the Victoria Stroop Test. *Aging*, Neuropsychology, and Cognition, 13(1), 20–35.

van der Kant, R., Goldstein, L. S., & Ossenkoppele, R. (2020). Amyloid-β-independent regulators of tau pathology in Alzheimer disease. Nature Reviews Neuroscience, 21(1), 21–35.

Warrington, E. K. (1984). Recognition memory test. NFER-Nelson.

Warrington, E. K. (1997). The graded naming test: a restandardisation. Neuropsychological Rehabilitation, 7(2), 143–146.

Wechsler, D. (2000). Wechsler Adult Intelligence Scale, third edition. Pearson Education.

Wilson, D., Le Heron, C., & Anderson, T. (2021). Corticobasal syndrome: a practical guide. Practical Neurology, 21(4), 276–285.

Wobbrock, J. O., Findlater, L., Gergle, D., & Higgins, J. J. (2011, May). The aligned rank transform for nonparametric factorial analyses using only anova procedures. In Proceedings of the SIGCHI conference on human factors in computing systems (pp. 143–146).

